# Lokatt: A hybrid DNA nanopore basecaller with an explicit duration hidden Markov model and a residual LSTM network

**DOI:** 10.1101/2022.07.13.499873

**Authors:** Xuechun Xu, Nayanika Bhalla, Patrik Ståhl, Joakim Jaldén

**Affiliations:** Division of Information Science and Engineering, KTH Royal Institute of Technology, Stockholm,11428, Stockholm, Sweden; Department of Gene Technology, Science for Life Laboratory, KTH Royal Institute of Technology, Solna, 17165, Stockholm, Sweden

**Keywords:** basecalling, HMM, LSTM, nanopore

## Abstract

**Background:** Basecalling long DNA sequences is a crucial step in nanopore-based DNA sequencing protocols. In recent years, the CTC-RNN model has become the leading basecalling model, supplanting preceding hidden Markov models (HMMs) that relied on pre-segmenting ion current measurements. However, the CTC-RNN model operates independently of prior biological and physical insights.

**Results:** We present a novel basecaller named **Lokatt**: exp<u>l</u>icit durati<u>o</u>n Mar<u>k</u>ov model and residu<u>a</u>l-LS<u>T</u>M ne<u>t</u>work. It leverages an explicit duration HMM (EDHMM) designed to model the nanopore sequencing processes. Trained on a newly generated library with methylation-free Ecoli samples and MinION R**9**.**4**.**1** chemistry, the Lokatt basecaller achieves basecalling performances with a median single read identity score of **0**.**930**, a genome coverage ratio of **99**.**750%**, on par with existing state-of-the-art structure when trained on the same datasets.

**Conclusion:** Our research underlines the potential of incorporating prior knowledge into the basecalling processes, particularly through integrating HMMs and recurrent neural networks. The Lokatt basecaller showcases the efficacy of a hybrid approach, emphasizing its capacity to achieve high-quality basecalling performance while accommodating the nuances of nanopore sequencing. These outcomes pave the way for advanced basecalling methodologies, with potential implications for enhancing the accuracy and efficiency of nanopore-based DNA sequencing protocols.

**Supplementary information:** Supplementary data are available online.

## 1 Introduction

The concept of nanopore-sequencing was first drafted in 1989 as a hand-sketched illustration by David Deamer on a page of a notebook [1]. 30 years later, the technology is now commercially available from Oxford Nanopore Technologies (ONT) [2]. Nanopore sequencing works by threading a single-stranded DNA (ssDNA) molecule through a protein-formed pore in a membrane, where the sequence of nucleotides along the ssDNA can be indirectly recorded through their effect on an ion current flowing through the pore. However, transforming the current measurements into a sequence of nucleotide bases, i.e., basecalling, is challenging for several reasons [3]. Firstly, multiple nucleotides along the ssDNA, also known as a *k*-mer where *k* is the number of nucleotides, simultaneously affect the noisy current measurements at any given time. Secondly, the ssDNA’s translocation speed through the pore is fast and unstable, leading to a random and apriori unknown number of current samples per nucleotide base in the measurements. Further, the short-term average translocation speed and the noise level are also variable across a long sequencing run due to measurement induced changes in the experimental conditions. These challenges made early nanopore sequencing unusable for most clinical and research applications. Although these problems have now been mitigated by, e.g., selecting and modifying various protein pores with narrower constriction, using improved DNA ratcheting enzymes to control the ssDNA’s translocation speed, and better basecalling algorithms, the technology is still limited for many applications due to high error rates, amounts of material, and costs [4–6].

Early basecalling algorithms worked by first segmenting the current measurements into a sequence of probabilistic events [7]. These events were then treated as observations in a graphical model, usually as a Hidden Markov model (HMM) with latent states representing the dominating *k*-mers [1, 8]. The representation of observation probabilities in HMMs went from simple Gaussian distributions parameterized by the mean and variance of the ionic current during an event [9, 10] to more elaborate models such as hierarchical Dirichlet processes [11]. With the probabilistic model in place, the final basecalling step could be completed using standard inference algorithms for HMMs, such as the Viterbi and the beam search (BS) algorithms. However, the performance of the HMM remained severely limited by the quality of the segmentation step and the choice of features used to model the distribution of the events.

To avoid the limitations of the HMM basecallers, most modern basecallers are based on end-to-end deep neural networks (DNN), following the pioneering work that led to the Chiron basecaller [12]. Specifically, Chiron applied a recurrent neural network (RNN) to process the current measurements and a connectionist temporal classification (CTC) layer from natural language processing (NLP) to replace the event-based HMM for data alignment. The resulting network was then trained in an end-to-end manner. The previously used event features have thus been replaced by neural networks, which obviate the need for explicit feature engineering, and the HMMs were replaced by the much simpler CTC structure. The ONT open-source software Bonito [13], which also uses a CTC-RNN structure and end-to-end training, performs on par with the current state-of-the-art commercial software Guppy, which presumably also uses a deep learning solution.

Inspired by these pioneering projects, recent research into basecallers has mainly explored variations in the neural network structure. Networks with recurrent units such as long-short term memory (LSTM) networks and RNNs, temporal convolution networks (TCNs), and attention/transformer networks such as the convolution-augmented transformer have all showed acceptable basecalling accuracy when combined with a CTC layer [14–16]. The attention structure could also be used without the CTC layer, which has shown superior performance in NLP applications [17]. Basecallers built solely with attention have, however, not yet demonstrated higher accuracy for basecalling [18]. This may be because nanopore signals have more vague transitions between nucleotides than words in NLP tasks, especially within homopolymers or repeated sequences of purines or pyrimidines.

This said, the lack of linguistic rules for DNA sequences does not mean there is no prior knowledge about the process that generated the nanopore data. For example, some studies model the ratcheting enzyme, e.g., a helicase, as finite state space machines with well-defined state transition probabilities driven by ATP concentration [19]. However, it is not straightforward to incorporate such knowledge in basecallers solely built with deep learning, although we believe that incorporating such prior knowledge could provide a pledge of in-depth understanding of nanopore sequencing and a new direction for future developments.

With this in mind, we wanted to revisit HMMs for basecalling while explicitly addressing some of the shortcomings of prior HMM basecallers. To this end, we propose a new hybrid basecaller called **Lokatt** that uses an explicit duration HMM (EDHMM) model with an additional duration state that models the dwell time of the dominating *k*-mer. The duration state allows the basecaller to be applied directly to the raw current measurements by circumventing the need to pre-segment the data, which was problematic in previous HMM basecallers. It also allows us to explicitly model the probability distribution of *k*-mer dwell times within the pore, e.g., based on a physical understanding of the ratcheting enzyme [19]. However, we still use a neural network to estimate the individual *k*-mer probabilities, and we train our basecaller using end-to-end training.

We trained and evaluated the Lokatt model on a methylation-free Ecoli dataset acquired locally with an ONT MinION device and Guppy. In order to establish a comparative baseline with the state-of-the-art architecture, we also trained the Bonito model from scratch using the identical training dataset. Additionally, to assess the models’ generalization capability, we extended our evaluation by employing a dataset from [20] that consists of ten different bacteria. Our benchmarking provides a proof of concept indicating that such hybrid models can exhibit comparable performance to the Bonito model in raw read accuracy and consensus assembly quality when training on the same dataset while opening up possibilities for engineered structures.

## 2 The Lokatt Model

An HMM is a generative Bayesian model used to represent the relationship between a sequence of latent states, which are assumed to be generated according to a Markov process, and a corresponding sequence of observations. For the basecalling task at hand we focus on building a hierarchical HMM structure to encode the temporal dependencies between a *k*-mer sequence of length *M*, denoted by ***K*** ≜ {*K*_1_, *K*_2_, …, *K*_*M*_} where *K*_*m*_ ∈ 𝕂 ≜ {1, …, 4^*k*^}, and a current measurement sequence of length *N*, denoted by {*X*_1_, *X*_1,2_, …, *X*_*N*_,}, where typically *M ≪ N*.

The duration of any state with a self-transition in a Bayesian state-space model is always geometrically distributed. This is inconsistent with the dwell times reported for both polymers and helicases [21, 22], two popular candidates for ratcheting enzymes. This inconsistency causes basecalling errors in regions rich in homopolymers since current variations are relatively small here, implying that the basecaller can only rely on the statistical modeling of the translocation speed. We address this problem via the introduction of a sequence of *M* explicit duration variables [23], denoted by ***D*** ≜ {*D*_1_, *D*_2_, …, *D*_*M*_} where *D*_*m*_ ∈ ℕ. This provides flexibility in terms of assigning an arbitrary dwell-time distribution. Each pair (*K*_*m*_, *D*_*m*_) for *m* = 1, …, *M* thus encodes the *m*th *k*-mer and its dwell time. However, to encode potentially very long dwell times using a limited number of states, we will also allow self-transitions between the states encoding the distribution of *D*_*m*_ for *m* = 1, …, *M*.

In the following two subsections, we formalize the hybrid data model on which Lokatt is based, beginning with the EDHMM and continuing with the DNN observation model.

### 2.1 The EDHMM structure

We consider a hierarchical and generative Bayesian EDHMM for the nanopore data, constructed as follows. We first draw the number of *k*-mers, *M* ∈ ℕ, from some distribution *ζ*(*M*). Then, we draw the first-order Markov sequence of *k*-mers ***K*** starting with *K*_1_ from a distribution *ξ*_0_(*K*_1_) followed by recursive draws of *K*_*m*_ from *K*_*m−*1_ according to a conditional distribution *ξ*(*K*_*m*_|*K*_*m−*1_) for *m* = 2, …, *M*. At the same time, we draw the sequence of dwell-times ***D*** by drawing each *D*_*m*_ independently from some distribution *η*(*D*) for *m* = 1, …, *M*. Finally, *D*_*m*_ measurements *X*(_*m,d*_) for *d* = 1, …, *D*_*m*_ are drawn for each *k*-mer *K*_*m*_ for *m* = 1, …, *M*, from a conditional distribution *φ*(*X*(_*m,d*)_ | *K*_*m*_). Here the state variable *d* acts as a counter that counts down to the next draw of a *k*-mer.

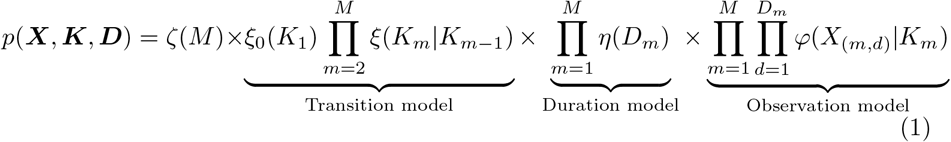

For notational convenience, we let ***X*** denote the sequence of measurements in the order of which they would be obtained, i.e.,

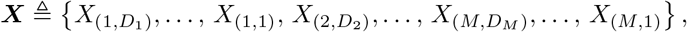

and use simply *X*_*n*_ for *n* = 1, …, *N*, where 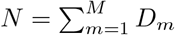, when speaking of the *n*th consecutive measurement. The joint probability distribution of the model is given in Eq. (1).

To allow for computationally efficient inference using the model, we make two additional assumptions on the model: First, we assume that the sequence length is geometrically distributed such that *ζ*(*M*) = (1− *α*)*α*^*M*^ for some *α* ∈ [0, 1); Second, we assume the duration distribution has a geometric tail such that *η*(*D* + 1) = *γη*(*D*) for all 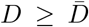 for some *γ* ∈ [0, 1) and some maximum explicit duration constant 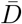 ∈ ℕ. The value of these assumptions is that they are inherently encoded in a *finite* state-space model using self-transitions between states. The generative model can, with these assumptions, be encoded as a state model with a start state *A*, an end state *B*, and 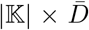 intermediate state pairs (*K, d*) representing joint *k*-mer and duration states. The intermediate states are stochastically reset according to the explicit duration probability *η* upon drawing new *k*-mers. The set of paths from *A* to *B* is isomorphic with the set of pairs (***K, D***), and a pair can be drawn from *p*(***K, D***) by a random walk from *A* to *B* with the following state transition rules:

1. State *A* transitions into state (*K*_1_, *d*) with probability *αξ*_0_(*K*_1_)*η*(*D* = *d*) for any 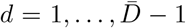, into state (*K*_1_, 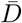) with probability 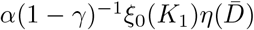, and directly into state *B* with probability 1− *α*.
2. State (*K*_*m*_, *d*) deterministically transitions into state (*K*_*m*_, *d*− 1) with the same *k*-mer for any *K*_*m*_ ∈ 𝕂, *d* = 2, …, 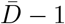, and *m*≥1.
3. State (*K*_*m*_, 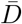) self transition into state (*K*_*m*_, 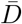) with the same *k*-mer with probability *γ* for any *K*_*m*_ ∈ 𝕂, and transitions into state (*K*_*m*_, 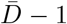) with probability 1− *γ*.
4. State (*K*_*m*−1_, 1) transitions into state (*K*_*m*_, *d*) with new drawn *k*-mer with probability *αξ*(*K*_*m*_|*K*_*m*−1_)*η*(*d*) for any (*K*_*m−*1_, *K*_*m*_) ∈ 𝕂 *×* 𝕂 and 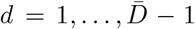, transitions into state (*K*_*m*_, 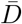) with probability 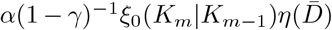, and transitions into state *B* with probability 1− *α*.

The state-space model is a standard implementation of an EDHMM [23]. The *m*th time the process returns to a state of the form (*K*_*m*_, 1) for *K*_*m*_ ∈ 𝕂 marks the end of *k*-mer *K*_*m*_ in the data. Each measurement, *X*_*n*_ = *X*(_*m,d*_), can also be drawn directly based on the value of state (*K*_*m*_, *d*), although we, in our particular implementation, assume that the observations are independent of *d*.

The value of the state-space representation is that it allows for efficient inference while maintaining model interpretability [24]. From the specific model presented above, the EDHMM can be constructed into a graph with size (*M×* 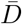) *× N*, as illustrated in Figure 1. The joint data likelihood and *k*-mer sequence *p*(**K, X**) can, using this graph, be efficiently computed with Eq. (1) by applying the forward algorithm [25, 26]. The data likelihood *p*(**X**) can, similarly, be efficiently computed by applying the forward algorithm to a slightly altered graph, where *K*_*m*_ ∈ 𝕂 can be arbitrary. The two graphs are sometimes referred to as the *clamped* and *free-running* models, respectively [25, 26], and allow us to efficiently compute the posterior sequence likelihood as *p*(**K X**) = *p*(**K, X**)*/p*(**X**). The conditionally most likely sequence of states and dwell-times (***K, D***) can be computed using the Viterbi algorithm, and the conditionally most likely sequence of states ***K*** can be approximately obtained using any of our recently introduced greedy marginalized BS (GMBS) algorithms [27].

**Fig. 1.**
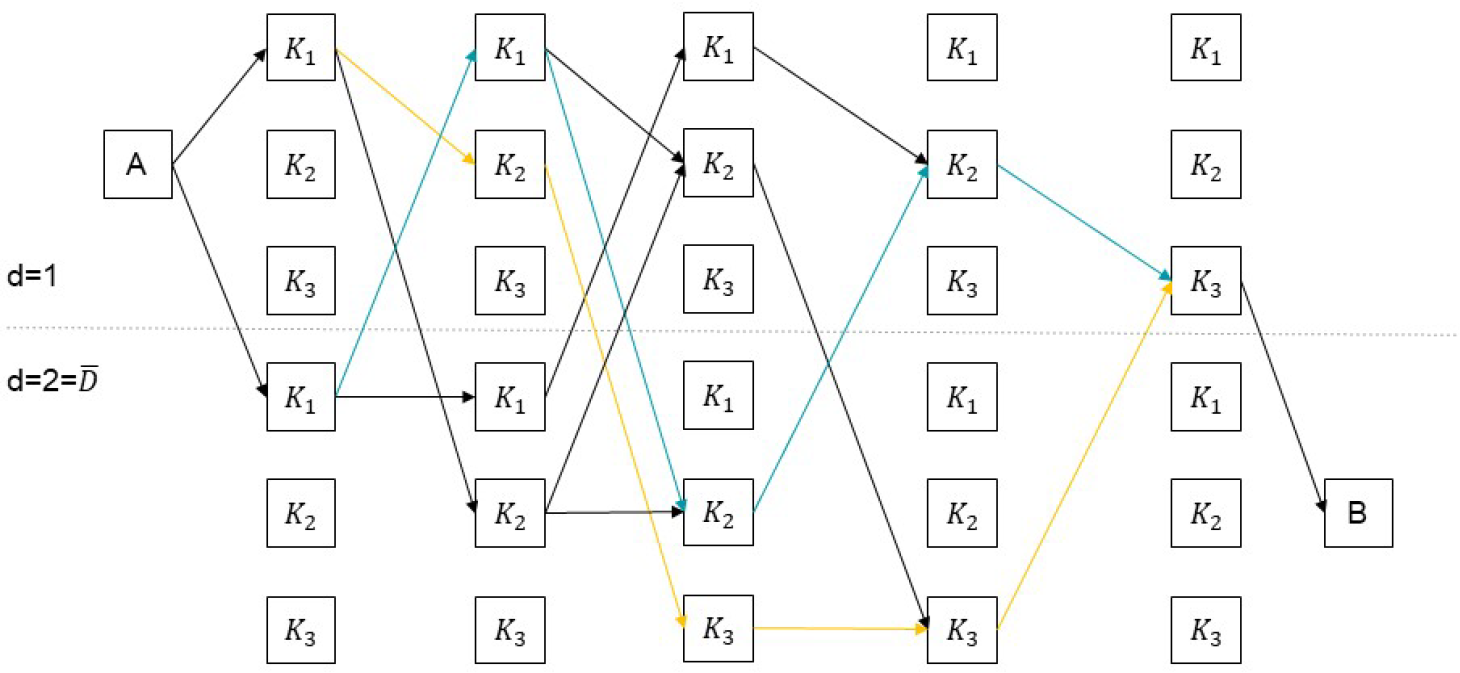
Illustration of EDHMM structure, with example of **K**_3_ = *{K*_1_, *K*_2_, *K*_3_*}* and **X**_5_ = *{X*_1_, …, *X*_5_*}*, hence *M* = 3, 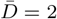 and *N* = 5. States in the upper half represent (*K*_*m*_, *d* = 1) while states in lower half represent (*K*_*m*_, *d* = 2). The blue path goes through five states (*K*_1_, *d* = 2), (*K*_1_, *d* =1), (*K*_2_, *d* = 2), (*K*_2_, *d* = 1), (*K*_3_, *d* = 1), representing the sequence of pairs (*K*_1_, *D*_1_ = 2), (*K*_2_, *D*_2_ = 2), (*K*_3_, *D*_3_ = 1); The yellow path goes through (*K*_1_, *d* = 1), (*K*_2_, *d* = 1), (*K*_3_, *d* = 2), (*K*_3_, *d* = 2), (*K*_3_, *d* = 1), representing the sequence of pairs (*K*_1_, *D*_1_ = 1), (*K*_2_, *D*_2_ = 1), (*K*_3_, *D*_3_ = 3).

Choosing the exact distributions in the model are also rather straightforward. The *k*-mer transition probabilities *ξ*(*K*_*m*_ | *K*_*m−*1_) can, for example, be estimated with a maximum likelihood (frequency counting) estimator from a reference genome; or by using uniform probabilities for *k*-mers *K*_*m*_ that originate from *K*_*m−*1_, e.g., set to 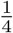 for each possible one-base transition and zero for any other combination, in order not to bias the model towards any particular genome. The value of *α* needs to be set very close to 1 for the model to plausibly generate reads on the order of hundreds of thousands of bases, and for all intents and purposes, one can set *α* = 1 in the implementation of the inference algorithms which we also do, although this does, strictly speaking, lead to an ill-defined prior distribution. The dwell-time distribution *η*(*D*_*m*_) can be estimated from sequences realigned at the sample-to-*k*-mer level. In the particular realization of Lokatt studied herein, we use a log-logistic distribution for *η* with parameters estimated using linear regression from the average number of large changes in the signal amplitude per sample. At the same time, we choose the tail factor *γ* so that *η*(*D*) provides a representative mean dwell time. We also perform a read-specific duration estimation during training and inference in our implementation since the average duration is longer for reads obtained later during the sequencing run due to ATP depletion. Finally, we replaced *φ*(*X*(_*m,d*_)|*K*_*m*_) with scores obtained from a match network 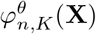, where *K* ∈ 𝕂, *n* = 1, …, *N* and where *θ* are the trainable weights of the (match) DNN described next.

### 2.2 The DNN structure

Lokatt relies on a neural network to dynamically extract the features from the current signal, then map them into scores associated with each *k*-mer, i.e., 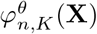. As we want the basecaller to be capable of handling various lengths of inputs, we construct a network with convolution kernels and recurrent units. In particular, we use two types of blocks: i) two residual-convolution blocks consisting of three convolution layers, with 32, 64, 32 features respectively, and a skip connection [28]; and ii) two bi-directional LSTM blocks [29], with 256 features in the first block on each direction and 1024 for the second block. In the end, a dense layer is applied to map the 2 *×* 1024 = 2048 features into |𝕂| = 4^*k*^ dimensions, which in Lokatt is 1024 with *k* = 5. Lokatt contains two of each block type, resulting in a total of 15.3 million weights. The complete network is shown in Figure 2 with a decomposition of the residual-convolution block and the bi-directional LSTM block. We used Swish activation, which nonlinearly interpolates between a linear function and the ReLU function [30, 31] between each layer in the network.

**Fig. 2.**
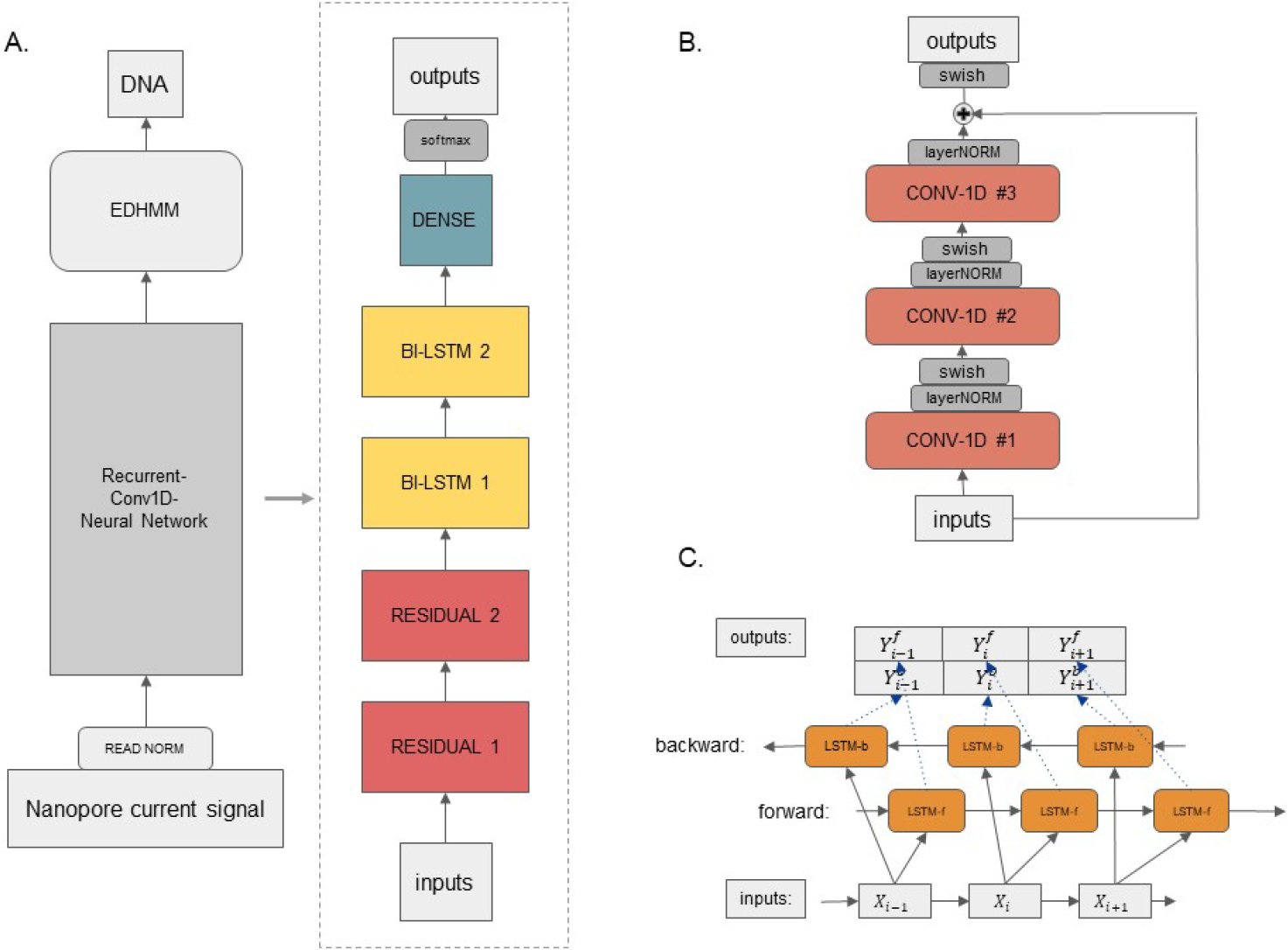
The overall structure of the Basecaller: **A**. The main components of Lokatt, from the bottom up, are normalized input, the neural network, the EDHMM layer, and output. The neural network is expanded into main component layer blocks on the right side: two residual blocks, two bi-directional LSTM clocks, and a dense layer. **B**. The inner structure of the residual block consists of three layers of 1D convolution, followed by layer-normalization, and the Swish activation, of which the outputs are taken and added with the inputs of the residual block followed by cross-layer normalization. **C**. The inner structure of the bi-directional LSTM layer consists of two independent LSTM layers, one in the forward direction and the other in the backward direction. The outputs of the two LSTM layers are then concatenated along the feature dimension, making the output sequence the same length as the inputs.

### 2.3 Model training

The weights in Lokatt’s DNN are trained end-to-end with (semi)-supervised learning using a modified conditional maximum likelihood (CML) approach. The vanilla CML objective maximizes the conditional log-likelihood, i.e. log *p*(**K** | **X**), which is often decomposed as log *p*(**K X**) =− log *p*(**K, X**) − log *p*(**X**). The clamped state-space model is used to compute log *p*(**K, X**) and the free-running model is used to compute log *p*(**X**), respectively, the former with latent states representing the particular target *k*-mer sequence and the latter with latent state from 𝕂 representing all possible sequences. Through the two graphs, the gradients can be computed with respect to 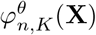and passed down to update the weights in the DNN with back-propagation [26].

During the training of Lokatt, we realized that this strategy led to a network that did not generalize well and gave low final scores. In a sense, the model learned to maximize log *p*(**K** | **X**) by minimizing log *p*(**X**) and hence did not provide a very good model of the data. This leads the neural network to output low values of the score 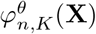 overall. Therefore, we instead adopted a modified CML training strategy, where we maximized a weighted linear combination of log *p*(**K** | **X**) and log *p*(**X**) to regularize the model to provide a balance between posterior decisions and modeling of the data. In particular, we explicitly used

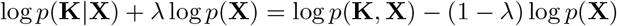

with regularization factor 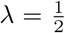 as the overall cost function. We also trained the network with a cost function where *p*(**X**) was replaced by *p*(**K**_BS_, **X**), i.e, with

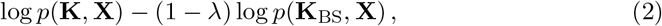

again with 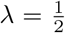, but where **K**_BS_ is the output sequence of *k*-mers from the GMBS decoder presented in Section 3.3 and discussed in detail in [27]. The intuition behind Eq. (2) is to make the training focus more on segments of the data where **K**_BS_ differ from the (correct) reference sequence **K**. The model trained with this approach showed higher final basecalling accuracy. Therefore, it was used to benchmark Lokatt with the other basecallers.

## 3 Methods

### 3.1 Data generation

To evaluate Lokatt, we performed ONT MinION sequencing on non-methylated *E*.*coli* genomic DNA (D5016, Zymo Research) in two repeated runs. The choice of this DNA type stemmed from an initial assumption regarding the potential significance of DNA methylation in basecalling. The sequencing libraries were prepared by fragmenting the genomic DNA using Covaris g-TUBE and a Ligation sequencing kit (SQK-LSK109, Oxford Nanopore). The first sequencing run, conducted in December 2019, used Flow Cell chemistry R9.4.1 and was basecalled with Guppy 3.2.2. The second sequencing run took place in November 2021 with Flow Cell chemistry R9.4.1 and Guppy 5.0, aimed at obtaining a recent comparison with state-of-the-art software and generating distinct datasets for training and evaluation purposes.

Furthermore, short-read Illumina sequencing was performed using TruSeq PCR-free library preparation on the MiSeq sequencing platform (Illumina, USA). A draft assembly was constructed from the Illumina data using SPAdes v3.6.0 [32], serving as the reference genome. The ground-truth nucleotide sequence for each raw read was obtained by mapping its tentative sequence, provided by Guppy, to the reference genome using the aligner program `Minimap2`[33].

The data were divided into batches based on their position on the genome. We divided the data from the 2019 sequencing run into batch 1.A and batch 1.B, by separating reads corresponding to the first and second half of the reference genome, respectively. The 2021 sequencing run was similarly divided into batch 2.A and batch 2.B. The data in batch 1.A was used as training data for the model, while the remaining batches were used for validation and performance evaluations. This allowed us to ensure that the model was not over-fitted to the underlying genome by comparing the performance on batch 2.A, where an obvious homology exists, with the performance on batch 1.B and 2.B where no clear homology between the training and test data should exist. Similar data division strategies have been used for human genome data sets, where the training and testing were based on different chromosomes [18].

For further assessment of the decoding efficacy and model generalization capabilities, we employed the test dataset established in [20], referred to as data batch 3. This dataset encompasses a diverse spectrum of microbial species, including four distinct variants of Klebsiella pneumoniae, along with six other bacterial entities, namely Acinetobacter pittii, Haemophilus haemolyticus, Serratia marcescens, Shigella sonnei, Staphylococcus aureus and Stenotrophomonas maltophilia. Notably, these samples comprise methylated sequences and underwent sequencing utilizing both R9.4 and R9.4.1 chemistries during the temporal interval spanning the years 2017 to 2018.

### 3.2 End-to-end training

Before end-to-end training, we pre-trained the DNN structure on a subset of re-aligned training data to expedite the subsequent computationally intensive modified CML training. Alignment was achieved using the EDHMM structure of the Lokatt basecaller, of which the observation probability table is obtained from the Baum-Welch algorithm [25, 26]. This alignment step could also be facilitated using the complete Lokatt model, which inherently incorporates alignment information. Based on the alignment, we segmented the long raw reads into 4096-sample segments aligned with reference sequences to enable GPU-based parallel computation. The DNN was then pre-trained to predict *k*-mer probabilities per sample using the cross-entropy loss. The pre-training on one epoch of the re-aligned data took 1.5 GPU hour.

As described in section 2.3, training with the regularized CML loss from Eq. (2) requires two graphs representing *p*(**K, X**) and *p*(**K**_BS_, **X**) to compute the gradients with respect to the output of the DNN, i.e., 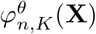, for *n* = 1, …, *N*. Note here that while *p*(**X**) is usually computed using the free running mode, we use the clamped model for both *p*(**K, X**) and *p*(**K**_BS_, **X**). In our implementation, the size of the two graphs are 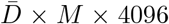 and 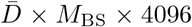, where the lengths of **K** and **K**_BS_, i.e., *M* and *M*_BS_, are on the order of a few hundred bases of overlapping *k*-mers. To manage the complexity of the inference on these graphs, we rely on custom GPU implementations that can efficiently utilize the sparsity of the graphs. In particular, we implemented the gradient calculation for the EDHMM as custom CUDA kernels [34] and registered them as customized operations in Tensorflow2 [35]. The gradients were then back-propagated to the DNN to calculate weight updates using a standard batch-based Adam optimizer [36] from Tensorflow. We trained the whole DNN on the NSC Berzelius compute cluster using 8 Nvidia A100 GPUs ^1^, for which training on one epoch of data took 576 GPU hours. The CUDA kernel code is available in the Lokatt repository: https://github.com/chunxxc/lokatt.

### 3.3 Decoding with the GMBS

In the final stage of basecalling, often referred to as *decoding*, the objective is to find the sequence of *k*-mer with the highest posterior probability, i.e. **K** = arg max_**K**_ *p*(**K** | **X**). The optimal solution to this problem is intractable due to the exponential growth of the search space as read length increases. This necessitates an exhaustive search across all possible *k*-mer sequences with lengths *M* that not exceeds *N*. An alternative involves utilizing the Viterbi algorithm, noted for its computational efficiency, to approximate the optimal solution by identifying the *jointly* optimal sequence (**K, D**) that maximizes *p*(**K, D**| **X**). Nonetheless, this approach lacks a solid theoretical guarantee regarding estimation quality. Moreover, the accurate decoding of the optimal *k*-mer sequence requires the marginalization of duration states on the EDHMM graph, a challenge for which the Viterbi algorithm lacks a dedicated strategy.

We instead rely on our newly proposed GMBS algorithms [27], which approximates the maximum a posteriori solution by recursively searching and maintaining a fixedsize list of *k*-mer sub-sequences **K**_*m*_ ≜ (*K*_1_, …, *K*_*m*_), i.e., *beams*, with high posterior probability *p*(**K**_*m*_|**X**_*n*_) for **X**_*n*_ ≜ (*X*_1_, …, *X*_*n*_) and *m ≤ n*. Each beam also keeps track of the probability of *p*(**K**_*m*_, *D*_*m*_|*X*_*n*_) from which it can compute the probability of new beams and marginalize over *D*_*m*_ as needed. The empirical experience in [27] has shown that the GMBS algorithm performs better than the Viterbi algorithm on decoding the EDHMM graph when we use 512 beams, with a significantly lower memory footprint. The GMBS performances can be improved by increasing the number of beams at the cost of slower pruning operations due to the sorting complexity increase. Specifically, the parallel sorting algorithms implemented in the GMBS scale, for a beam list size of *B*, as 𝒪 (*log*^2^*B*) [37].

In implementing the GMBS algorithm, we use a tree structure to store **K**_*m*_ in memory, where common ancestry represents common initial sub-strings of **K**_*m*_. Since this tree has at most *N× B* nodes, it can be efficiently implemented on the GPUs without needing dynamic memory allocation. Finally, to extract the final approximate maximum a posteriori *k*-mer sequence **K** = **K**_*M*_, we can read this tree backwards from the highest-scoring leaf node to the root in a fashion similar to the backtracking step of the Viterbi algorithm. A detailed explanation of the LFBS algorithm is provided in [27].

When basecalling with Lokatt, we divide each raw read into segments of length 4096 with an overlap of 296 measurements, and these segments were individually basecalled. The uniform length of input sequences facilitates efficient parallel implementations of Lokatt without harming the basecalling performance. The resulting output sequences were subsequently assembled by aligning the beginning and ending portions of consecutive pairs.

## 4 Results

### 4.1 Benchmarking

We benchmark the Lokatt model with the Bonito model over all three data batches. Bonito [13] is an open-source research tool released by ONT that harnesses the state-of-the-art CTC-RNN model. To investigate the impact of the model architecture, we independently trained a Bonito model, referred to as Bonito Local, from scratch with the same training data used in training the Lokatt basecaller. Additionally, we included the performance from the fine-tuned Bonito basecaller *dna_r*9.4.1_*e*8_*sup*@*v*3.3, denoted as ‘Bonito Sup’, which yields to give ‘super accuracy’ and is presumably trained on a more extensive data set. For data batches 2.A and 2.B, we also incorporated results obtained from ONT’s Guppy 5.0, which is ONT’s commercial basecaller. The data batches 1.A and 1.B were sequenced with earlier version of Guppy 3.2.2, whose performance is 2% − 5% lower than Guppy 5.0 and therefore excluded in the benchmark as it no longer represents state-of-the-art. In the following sections, we will discuss the basecalling quality in terms of raw read accuracy and assembly accuracy.

### 4.2 Raw read accuracy

To access the raw read accuracy, we measured the shortest Levenshtein distance between the basecalled sequences from Lokatt, Bonito Local and Bonito Sup and the reference genome and generated their pair-wise alignments. This entails determining the minimal number of single-state edits, such as insertions, deletions or substitutions, needed to convert the basecalled sequence into the Illumina-generated reference genome [38], which can be obtained using Minimap2 v.2.17-r941 with the default parameters. The alignments are then quantified and reported as the sequence accuracy metrics, including the identity, mismatch, insertion and substitution scores. Specifically, the identity score is formulated as the ratio of matched bases to the total alignment columns as follows:

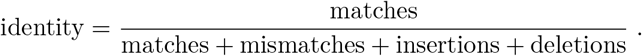

Similarly, the error scores, including mismatch, insertion and substitution, are calculated as the ratios of their individual count to the overall alignment columns. In addition, we also presented measurements of matched base length per read (read length), counts of matched read entries (entry counts) and the cumulative number of matched bases. It is important to note that both mean and median values were reported for accuracy metrics and read length. However, for our analysis, the focus was directed towards median values. This choice is attributed to the significant data variance, wherein reads with low accuracy are too noisy for Minimap2 to recognize [7][20].

The outcomes of the benchmarking experiments are presented in Tables 1 to 5, each highlighting the basecaller performances across different data batches. Table. 1 displays the training performance on data batch 1.A. Lokatt achieves a median identity score of 0.926, surpassing Bonito Local by 0.012 but falling 0.028 behind Bonito Sup, with corresponding lower error rates than Bonito Local and higher error rates than Bonito Sup. Regarding median read length, Lokatt records the longest matched length of 2807, compared to both Bonito models’ 2731. However, for recognizable read entries based on Minimap2, Lokatt processes 565393 entries. This number is greater than Bonito Local’s 562838, yet less than Bonito Sup’s 583471. Consequently, Lokatt demonstrates 3% fewer total matched bases than Bonito Sup, but 3% more than Bonito Local.

**Table 1.** Training performances on data batch 1.A.

| Basecaller | identity | insertion | deletions | substitution | base per read | entry counts | bases ($10^9$) |
| --- | --- | --- | --- | --- | --- | --- | --- |
| Lokatt | 0.912/0.926 | 0.018/0.016 | 0.045/0.034 | 0.025/0.020 | 4803/2937 | 565393 | 2.600 |
| Bonito Local | 0.900/0.914 | 0.019/0.017 | 0.051/0.038 | 0.030/0.026 | 4724/2885 | 562838 | 2.524 |
| Bonito Sup | 0.936/0.954 | 0.013/0.010 | 0.030/0.018 | 0.021/0.015 | 4757/2829 | 583471 | 2.686 |

\*first five entries are listed as ‘mean/median’.

Table 2 provides the test performance of data batch 1.B, which corresponds to reads aligned with the second half of the E.coli genome. The results indicate the median identity scores are within *±* 0.001 difference from the training scores shown in Table 1, with error rates exhibiting a similar *±* 0.002 difference. This minimal variation between batch 1.A and 1.B suggests that overfitting of the models is unlikely. Notably, Lokatt continues to exhibit the longest read length, while Bonito Sup maintains the highest entry count. This also contributes to Lokatt reporting 3% fewer matched bases than Bonito Sup, but 3% more than Bonito Local.

**Table 2.** Testing performances on data batch 1.B.

| Basecaller | identity | insertion | deletions | substitution | read length | entry counts | bases ($10^9$) |
| --- | --- | --- | --- | --- | --- | --- | --- |
| Lokatt | 0.912/0.926 | 0.018/0.016 | 0.046/0.035 | 0.024/0.020 | 4852/2961 | 523191 | 2.432 |
| Bonito Local | 0.899/0.913 | 0.020/0.018 | 0.052/0.040 | 0.030/0.026 | 4770/2917 | 521186 | 2.362 |
| Bonito Sup | 0.937/0.955 | 0.012/0.009 | 0.031/0.018 | 0.020/0.015 | 4799/2916 | 540834 | 2.513 |

\*first five entries are listed as ‘mean/median’.

Tables 3 and 4 display the testing result from data batch 2.A and 2.B, which were generated from the same E.coli samples used in data 1 but more recently. Compared to results from data batch 1.A and 1.B, all basecaller exhibit identity score 0.007–0.014 higher and, at most, error score 0.008 lower. Regarding the read length, Bonito Local shows an increase of 5%, while both Lokatt and Bonito Sup demonstrate an increase of 14% and 13%, respectively. The substantial increase in entry counts and total matched bases is because data batch 2 sequenced 3–4 times more samples, which makes the comparison trivial. Nonetheless, when comparing among basecallers on data batch 2, Lokatt demonstrates identity scores 0.006 and 0.007 higher than Bonito Local data 2.A and 2.B respectively and 0.033 lower than Bonito Sup, while output 5% fewer matched bases than Bonito Sup with 7% more than Bonito Local in total.

**Table 3.** Testing performances on data batch 2.A.

| Basecaller | identity | insertion | deletions | substitution | read length | entry counts | bases ($10^9$) |
| --- | --- | --- | --- | --- | --- | --- | --- |
| Lokatt | 0.903/0.933 | 0.016/0.013 | 0.051/0.033 | 0.029/0.020 | 5600/3398 | 1519444 | 8.132 |
| Bonito Local | 0.903/0.927 | 0.016/0.014 | 0.050/0.033 | 0.030/0.024 | 5392/3063 | 1467918 | 7.556 |
| Bonito Sup | 0.929/0.966 | 0.013/0.008 | 0.033/0.013 | 0.024/0.013 | 5588/3304 | 1579899 | 8.522 |
| Guppy 5.0 | 0.907/0.936 | 0.019/0.015 | 0.042/0.027 | 0.031/0.021 | 5637/3391 | 1522808 | 8.153 |

\*first five entries are listed as ‘mean/median’.

**Table 4.** Testing performances on data batch 2.B.

| Basecaller | identity | insertion | deletions | substitution | read length | entry counts | bases ($10^9$) |
| --- | --- | --- | --- | --- | --- | --- | --- |
| Lokatt | 0.904/0.933 | 0.016/0.013 | 0.052/0.034 | 0.028/0.020 | 5685/3494 | 1400186 | 7.612 |
| Bonito Local | 0.902/0.926 | 0.016/0.014 | 0.052/0.034 | 0.030/0.024 | 5473/3124 | 1353165 | 7.073 |
| Bonito Sup | 0.930/0.966 | 0.013/0.007 | 0.034/0.013 | 0.023/0.013 | 5660/3371 | 1459829 | 7.980 |
| Guppy 5.0 | 0.907/0.935 | 0.020/0.015 | 0.043/0.027 | 0.030/0.021 | 5720/3480 | 1403826 | 7.631 |

\*first five entries are listed as ‘mean/median’.

Additional results from Guppy 5.0, ONT’s commercial software at the experiment’s time, are also reported in Tables 3 and 4. Guppy’s overall performance closely aligns with Lokatt’s; Guppy exhibits a 0.003 higher identity score and basecalled only 0.25% more total bases.

Table 5 presents the testing performance on data batch 3 with diverse species sequenced at an earlier period. Table 5 only contains identity score and total base counts due to space limit; however, a comprehensive performance table is available in Table S1, Additional file 1. Notably, all basecaller exhibit reduced identity scores for all species except Staphylococcus aureus, most recently sequenced among data 3 and utilizing the updated chemistry R9.4.1. Overall, Lokatt demonstrates an average identity score of 0.904, which declined by 0.029 on average and 0.049 on maximum, compared with data 2; Bonito Local shows an average identity score of 0.880, with a decline of 0.046 and maximum decline of 0.079; Bonito Sup has an average 0.947 with a decline of 0.019 on average and 0.031 in maximum. For total matched bases, Lokatt holds 6% more than Bonito Local but 7% fewer than Bonito Sup and 0.3% fewer than Guppy.

**Table 5.**
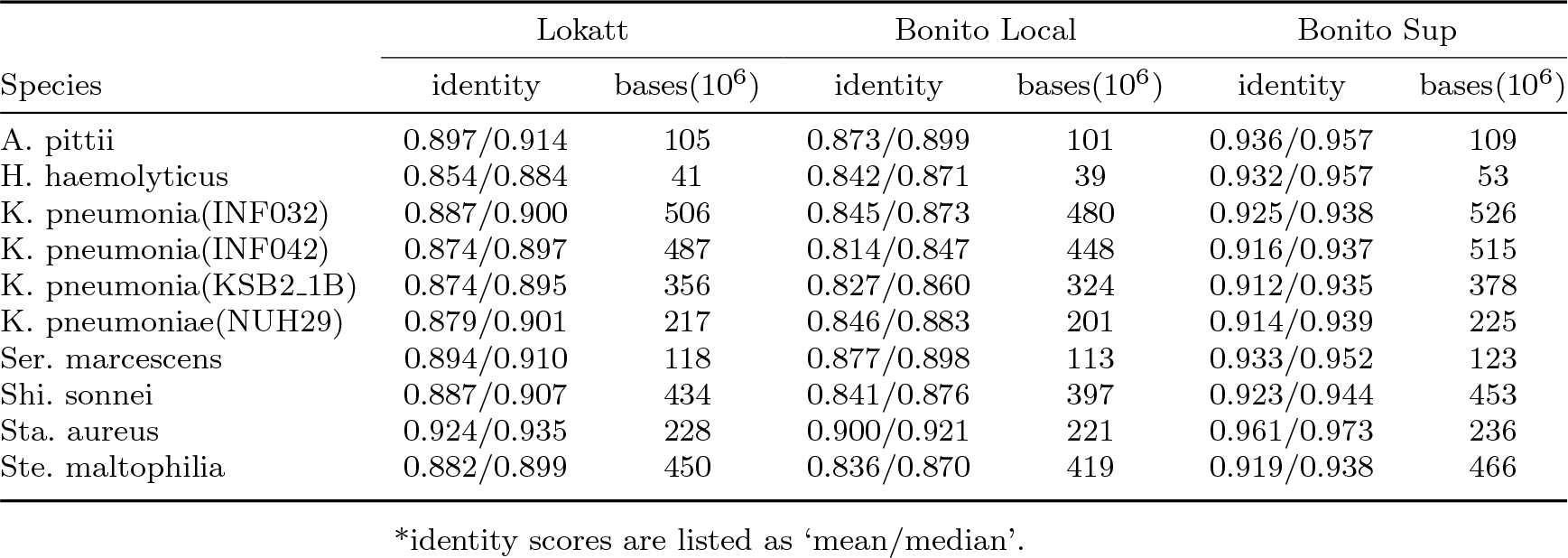
Testing performances on data batch 3.

### 4.3 Basecaller complexity

Incorporating a DNN structure, both Lokatt and Bonito introduce computational demands that can potentially restrict their applicability. Bonito 0.5.0 utilizes the CTC-Conditional Random Field [39] top layer, integrated with convolution layers followed by LSTM layers in alternating forward and reverse directions, totalling 26.9 million parameters. In contrast, Lokatt adopts residual blocks comprising convolution layers, followed by bi-directional LSTM layers and an EDHMM layer on top, collectively having 15.3 million parameters. It is noteworthy that Bonito’s employment of a stride of 5 in its convolution layer has effectively reduced the computationally expensive recurrent calculations within the LSTM layer by a factor of 5. Consequently, despite being nearly twice large as Lokatt, Bonito (50k base pair per second) achieves approximately a 5-fold speed enhancement compared to Lokatt (8k base pair per second) when executed on the Nvidia V100 GPU.

### 4.4 Consensus evaluation

The basecalled read sequences on data batch 2 from Lokatt, Bonito Local, Bonito Sup and Guppy 5.0 were assembled, respectively. De novo genome assemblies were generated using Flye [40], resulting in four distinct draft genome assemblies. The evaluation of these assemblies against the reference Illumina *E*.*coli* genome was executed with Quast [41]. The assessment, as depicted in Figure 3a, reveals a genome coverage of 99.750% for Lokatt, while the remaining three basecallers exhibit a slightly higher coverage of 99.757%. The proportions of GC content remain comparably consistent among the four basecallers: Lokatt at 50.88%, Bonito Local at 50.84%, Bonito Sup at 50.82%, and Guppy 5.0 at 50.83%, in contrast to the reference genome’s 50.77%, as illustrated in Figure 3b. The same contig length distribution and contig connectivity are shared among all basecaller’s assemblies, reflected in the NGAx plot in Figure 3c. The NGA50 values, specified in Figure 3d, exhibit marginal disparities: 106101 for Lokatt, 106277 for Bonito Local, 106360 for Bonito Sup, and 106293 for Guppy 5.0. Notably, Lokatt shows the least number of misassemblies at 202, compared to 204 for both Bonito basecallers and 203 from Guppy, as depicted in Figure 3e. Furthermore, the assessment of mismatches per 100k base pairs, illustrated in Figure 3f, highlights Lokatt’s score of 8.22k, which is notably lower than Bonito Local’s 9.9 and Bonito Sup’s 9.69, and approaches the minimum value of 8.14 attained by Guppy. The complete assemblies report is available in Table S2, Additional file 1.

**Fig. 3.**
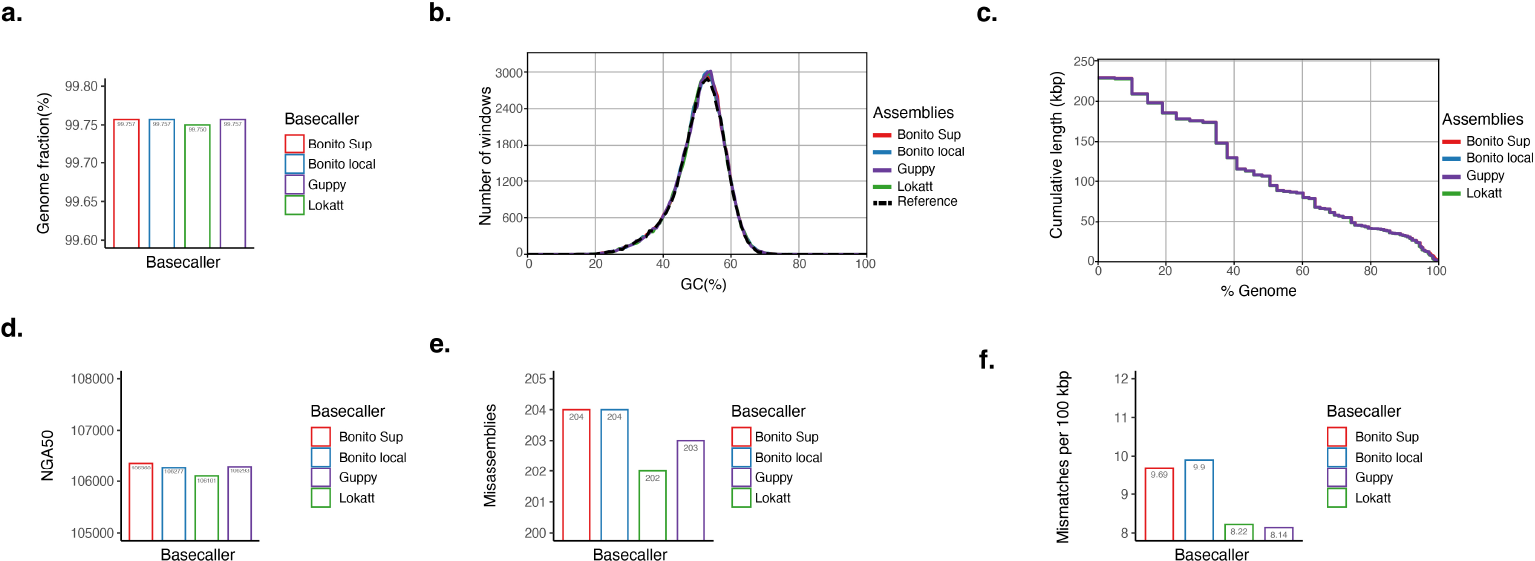
Performance plots for basecallers: A. Genome coverage ratios. B. The number of windows for each GC percentage. C. NGAx plots for the different assemblies showing the aligned contig length distribution against the reference genome. D. The NGA50 plot. E. The misassemblies. F. The mismatches per 100 kbp.

## 5 Discussion

The evaluation of basecallers, Lokatt, Bonito Local, Bonito Sup, and Guppy 5.0, on various datasets yielded insights into their performances and limitations. Analyzing raw read accuracy in E.coli datasets 1 and 2, Lokatt’s performance emerged as competitive with the Bonito basecallers. Lokatt achieved a slightly higher median identity score than Bonito Local by 0.01, while falling slightly behind Bonito Sup by 0.033. The assessment of generalization capabilities on the extra data batch 3 showed a decrease in performance for both Lokatt and Bonito Local, with the latter experiencing a more notable decline. This drop could be attributed to a mixture of the samples covering various genomes, as well as using old chemistry at earlier dates since Bonito Sup also showed a performance drop. Nevertheless, Bonito Sup exhibited superior accuracy, although limited information was available regarding its training data and methodology. Notably, the study underscored the Lokatt model’s capability to effectively handle small-genome species sequencing data at a level comparable to the Bonito model.

The assembly analysis introduced a noteworthy observation: the correlation between genome coverage and raw read accuracy across basecallers isn’t always straightforward. Specifically, Lokatt, despite having cumulatively 7% more matched bases than Bonito Local and a close number to Guppy, exhibited marginally 0.007% lower genome coverage on the E.coli genome. However, Lokatt maintained the least misassemblies and relatively low mismatches per 100kbp. Bonito Local on the other hand, having the lowest identity score and fewest base counts, achieves the same genome coverage as Bonito Sup, which shares an identical model architecture with Bonito Local but with different weights. This discrepancy might arise from differences in error-handling strategies among basecallers, potentially indicating that Lokatt excels in specific genomic regions but faces challenges in others. Alternatively, this could be attributed to Lokatt’s current lack of quality scores associated with basecalled nucleotides, which play a crucial role in the assembly process and subsequent analyses. Furthermore, it’s important to note that the evaluation primarily focused on E.coli data, which may not fully capture the challenges posed by diverse species and complex genomic regions.

The study also highlighted the distinct design and training strategies of Lokatt. Bonito utilizes the CTC loss, where a blank state is manually inserted between every nucleotide. The CTC loss typically leads to a dominance of blank predictions in the output of the CTC-trained RNN [12], prompts presumably a less complicated task where the RNN has learned to detect the transition moment of the nucleotides and treat everything else as blank. However, it also potentially limits model flexibility. In contrast, Lokatt integrates an EDHMM modelling the dynamic of the ratcheting enzyme, and is tasked to learn the complete characteristics of the ion current measurements. We have separated the complicated task into the pre-trained stage and the subsequent end-to-end training. We observed that the pre-trained neural network could achieve a median basecalling accuracy of around 0.905 on data batches 1 and 2. The subsequent CML training further refined Lokatt’s accuracy by around 0.02**–**0.03. In light of this, although HMMs are capable and comparable in various respects, achieving fully interpretable parametric models remains a challenge, leading to limitations in Lokatt’s performance.

## 6 Conclusion

This research project has culminated in the development, training, and evaluation of Lokatt, a novel hybrid basecaller. The raw read performance of the Lokatt model was better than the CTC-CRF structured Bonito model if trained on the same dataset, and was comparable to ONT-trained Bonito and Guppy basecallers. Lokatt’s evaluation metrics for consensus sequencing resembled those of the other basecallers. Notably, Lokatt demonstrated fewer misassemblies and mismatches per 100kbp, despite having a lower overall genome coverage ratio. This scenario highlights the complex trade-offs that basecallers need to make between accuracy, coverage, and the nature of the underlying genomic sequences. Different basecallers may excel in different contexts, and the choice of which to use depends on the specific goals of an analysis, such as accurate assembly of specific regions versus comprehensive genome coverage.

Both Lokatt’s and Bonito’s architectures leverage DNN structures enhanced with dynamic models to bridge the gap between input current measurements and output bases and enable end-to-end training. However, Lokatt’s unique integration of an HMM layer introduces a novel dimension of comprehension into the sequencing dynamics, thereby creating opportunities for future refinements. Future versions of Lokatt could potentially exploit a more sophisticated dynamic structure, informed by a deeper understanding of the sequencing device and chemistry process. This study contributes to the field by introducing an innovative basecaller model and insights into basecaller performance.

## Declarations

### Ethics approval

Not applicable.

### Consent to participate

Not applicable.

### Consent for publication

Not applicable.

### Availability of data and materials

The datasets generated during and/or analysed during the current study, including fast5, fasta, fastq and the reference genome files, are held on Zenodo, DOI 10.5281/zenodo.7970715 available on https://zenodo.org/record/7995806.

### Code availability

The code of Lokatt is available at this GitHub repository: https://github.com/chunxxc/lokatt.

### Authors’ contributions

XX and JJ are the main contributors to the design of the work, analysis and interpretation of data, coding and writing and revising the paper. NB and PS generated the E.coli data library, contributed to the assembly experiment and analysis, and revised the paper. All authors read and approved the final manuscript.

## Acknowledgements

The authors acknowledge support from the National Genomics Infrastructure (NGI) in Stockholm funded by Science for Life Laboratory and the Knut and Alice Wallenberg Foundation for assistance with massively parallel sequencing. The model training of Lokatt and basecalling were enabled by the supercomputing resource Berzelius provided by National Supercomputer Centre at Linköping University and the Knut and Alice Wallenberg foundation. The assemblies of the basecaller datasets were enabled by resources provided by the National Academic Infrastructure for Supercomputing in Sweden (NAISS) and the Swedish National Infrastructure for Computing (SNIC) partially funded by the Swedish Research Council through grant agreement no. 2022-06725 and no. 2018-05973. Finally, the authors wish to thank Josephine Sullivan for helpful discussions regarding the dimensioning and training of the neural networks used in Lokatt. We would also like to thank Carl Rubin and Remi-André Olsen for their input on nanopore sequencing and processing of data.

## Supplementary files

Additional file 1: An Excel file contains Table S1 Testing performances on data batch 3, including full sequence accuracy metrics of the benchmarking basecallers, and Table S2 Assemblies report on data batch 2, the comprehensive Quast genome assemblies report of all benchmark basecallers.

## Footnotes

1 Berzelius: https://www.nsc.liu.se/support/systems/berzelius-getting-started/

